# Tbx genes influence early gene expression and photoreceptor patterning in the chick retina

**DOI:** 10.1101/2025.10.18.683251

**Authors:** Monika Ayten, Heer N.V. Joisher, Constance Cepko

## Abstract

The retina, our sensory organ for vision, displays distinct photoreceptor distributions and can have a specialized region for high visual acuity. The establishment of these spatial patterns during development depends on regionally restricted transcription factors and signaling molecules. Members of the T-box family of transcription factors, Tbx2, Tbx3, and Tbx5, show such restricted patterns of expression, with a dorsal high to ventral low pattern in the early retina. Their potential regulatory interactions and roles in patterning have not been fully explored. Here, we investigated their regulatory interactions by overexpression (OE) and knockdown (KD) approaches. We found that Tbx2, Tbx3, and Tbx5, directly or indirectly regulate each other as well as other early, patterned genes, including *Fgf8, Cyp26a1, Cyp26c1, Cyp1b1, Raldh1, Ventroptin,* and *Bmp2*. KD of any one of these Tbx genes increased rod photoreceptor density in a specific region of the retina, whereas Tbx2 loss additionally reduced the number of UV cones throughout the retina. Notably, KD of Tbx2, Tbx3, or Tbx5 consistently resulted in a smaller rod-free zone (RFZ), a domain within the high acuity area. These findings demonstrate that T-box transcription factors form a coordinated regulatory network that governs regional gene expression and photoreceptor patterning.

**Highlights:**

- Tbx2, Tbx3, and Tbx5 regulate each other during chick retinal development
- Tbx2, Tbx3, and Tbx5 regulate the expression of early patterned genes, including *Fg8, Cyp26c1, Cyp26a1, Cyp1b1,* and *Bmp2*
- Knockdown of either Tbx2/3/5 increased rod numbers in the equatorial retina and reduced the size of the rod-free zone (RFZ)
- Tbx2 knockdown decreased UV cone numbers dorsally, ventrally and in the RFZ
- Tbx2, Tbx3, and Tbx5 had no impact on green or red cone abundance

## Introduction

Vision is initiated by the capture of light by highly specialized retinal cells, the photoreceptors. Among photoreceptors there is a wide range of sensitivity to light, with rod photoreceptors able to detect very few photons, enabling night vision (1). Cone photoreceptors are typically much less sensitive and initiate our daylight and color vision (2). Rods and cones are not equally distributed within the retina but exhibit patterns that can influence multiple aspects of vision, including acuity. Species with high acuity vision, like birds of prey and humans, have a specialized high acuity area (HAA) characterized by a high density of cone photoreceptors and the absence of rod photoreceptors. Moreover, different types of cones, defined by their morphology and tuning to different wavelengths of light, can also be patterned. For example, in humans, “red” and “green” cones are the only photoreceptor types in the central HAA, the foveola. Rods are excluded from this region (also known as the rod-free zone, RFZ) but are the most abundant photoreceptor type throughout the majority of the rest of the human retina, and in almost all mammals (3, 4). In addition to photoreceptors, other retinal cell types are distributed in specific patterns. The output neurons of the retina, the retinal ganglion cells, are most abundant in the central retina of many species. They also exhibit patterning throughout the retina, with their projections to the brain forming a topographic map of the visual field (5, 6).

The mechanisms that drive photoreceptor patterning have not been well elucidated. Our lab has been exploring this topic in the chicken, as we discovered that they have a central RFZ (7) in an area with other anatomical features of a HAA (8). The RFZ has a high density of cones with elongated outer segments, as in humans. There is also a higher density of retinal ganglion cells in this area, and an aster, a specialized structure in the inner nuclear layer of the retina (8, 9).The chick retina provides an excellent model to study photoreceptor patterning due to its accessibility early in development, when patterns are being established. We have used it as a model of HAA development and have started to identify some key regulators of the formation of this area. Lack of retinoic acid (RA) signaling and presence of Fgf8, both of which are specific to a circumscribed central area, are required for formation of the chick HAA (8). RA signaling gradients also have been implicated in cone subtype specification in other species (10). Ventrally restricted expression of cVax is required for the dorsal-ventral gradient of rods across the entire retina (11). We have further examined the expression of other genes that are expressed very early in development, with patterns suggestive of the regulation of photoreceptor pattern (Joisher et al, BioRxiv). These include *Fgf8, Cyp26c1, Cyp26a1, Raldh1, Cyp1b1, Bmp2,* and *Ventroptin.* The regulators of these patterned genes have not been identified, but transcription factors that are also expressed very early are potentially part of their regulatory network.

The T-box (Tbx) genes provide one such example of potential regulators. They encode a family of evolutionary conserved transcription factors that play essential roles in embryonic development and organogenesis (12, 13). During retinal development, *Tbx2, Tbx3,* and *Tbx5* are expressed in the dorsal optic cup of humans, mice, and chicks (14–16), where they exhibit distinct but partially overlapping expression domains. Tbx proteins are characterized by a conserved DNA-binding motif, or T-box domain, that binds DNA in a sequence-specific manner. Several Tbx genes display partially redundant functions and act as key regulators of cell fate decisions, tissue patterning, and differentiation across multiple organs, including the nervous system and sensory structures (12, 13).

The expression patterns of the Tbx genes suggest both unique and overlapping roles in shaping retinal patterns and cell fates (Joisher et al., BioRxiv). For example, *Tbx5* is known to determine dorsal identity and thereby contribute to the formation of the topographic map made by the projections of the retinal ganglion cells (15, 17). The *Tbx2* orthologue in zebrafish, Tbx2b, inhibits rods and promotes UV cone fate (18, 19). This makes the dorsally restricted expression of *Tbx2, Tbx3,* and *Tbx5* in the chick retina particularly intriguing, as the dorsal chick retina has a relatively low rod density (7). Moreover, a restricted central domain of Tbx3 expression overlaps with the developing rod-free HAA during early development (Joisher et al., BioRxiv). These observations raise the possibility that Tbx factors not only contribute to overall dorsoventral patterning but also participate in the formation of the RFZ within the HAA.

Here, we examined the expression patterns and functions of these Tbx genes in the developing chick retina. Specifically, we focused on how their activity influences dorsoventral patterning, early retinal gene expression, and the spatial distribution of photoreceptor subtypes, including the formation of the RFZ. Beyond informing our understanding of the development of this important area, these findings set a framework for understanding the interplay of early patterning events with later cell fate choices in an accessible system.

## Results

### Tbx genes exhibit distinct spatial expression patterns in the developing chick retina

To investigate the spatial expression patterns for *Tbx2, Tbx3,* and *Tbx5* during retinal development, we examined chick retinal flatmounts from embryonic day (E) 3 to E8 using the hybridization chain reaction (HCR) method of single molecule fluorescent *in situ* hybridization (smFISH) (Fig. 1A). This developmental window was selected because it encompasses key stages of retinal neurogenesis and lamination. Between E3 and E3.5, retinal progenitor cells begin to produce daughter cells that exit the cell cycle, with the first born cells, the retinal ganglion cells, differentiating in the central retina (20). Neurogenesis continues until it is nearly complete by E8 (21). Retinal lamination is ongoing during this period, with the sequential addition of horizontal, amacrine, and photoreceptor cells (22, 23).

**Figure 1:**
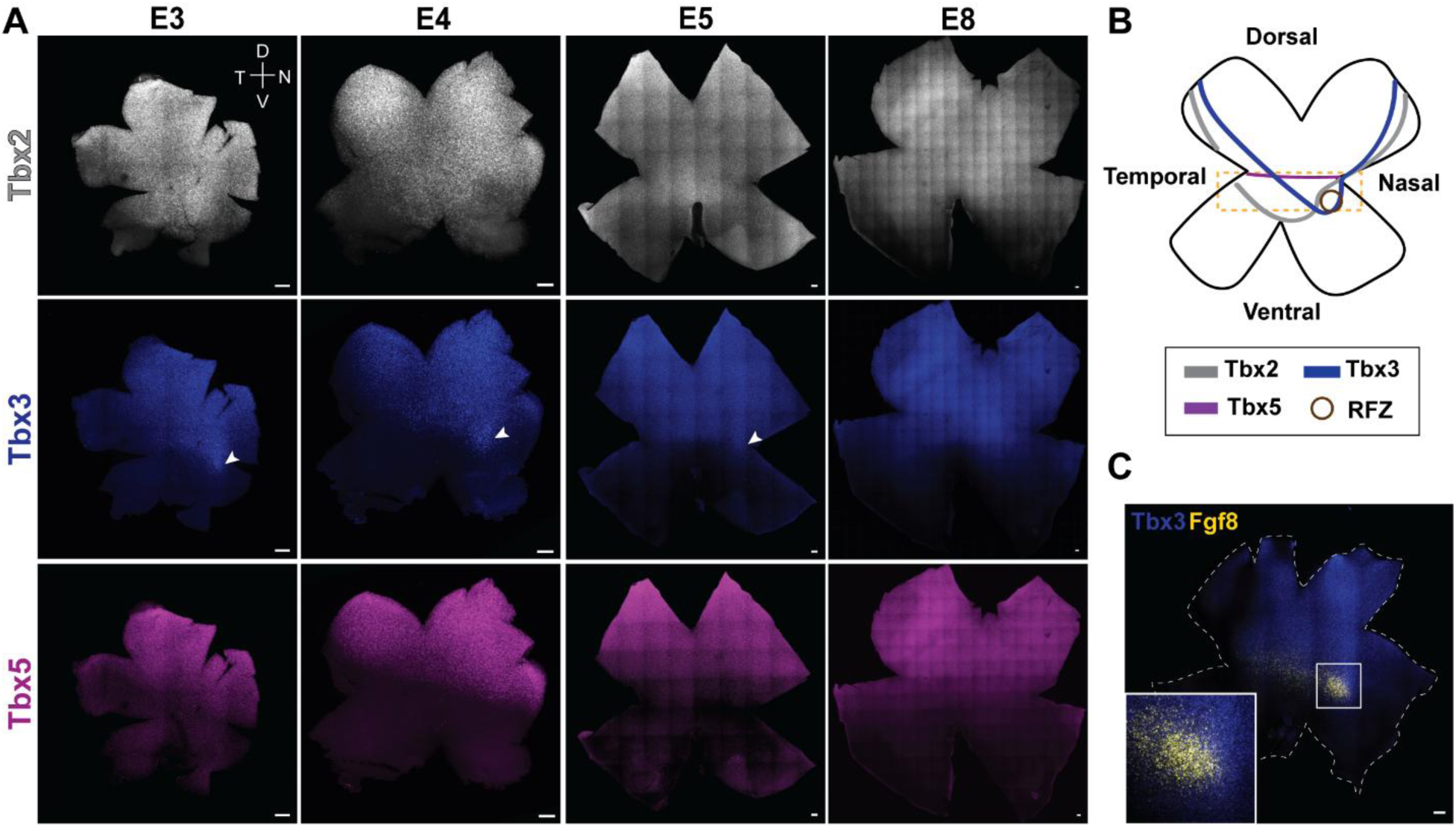
Spatial expression patterns of Tbx2, Tbx3, and Tbx5 in the developing chick retina. **(A)** Representative smFISH images of retinal flatmounts from embryonic day (E)3 to E8 probed for Tbx2, Tbx3, and Tbx5. The white arrowhead marks a spot of Tbx3 that overlaps with Fgf8 (and see panel C) **(B)** Schematic representation of the expression domains of Tbx2 (gray), Tbx3 (blue), and Tbx5 (magenta) in the retina. The yellow box indicates the equatorial retinal domain. **(C)** smFISH image of a retinal flatmount from E5, probed for Tbx3 and Fgf8. **(A,C)** Scale bar, 100 µm.

All three genes, *Tbx2, Tbx3,* and *Tbx5,* are expressed in the dorsal retina (Fig. 1A), with *Tbx2* extending furthest ventrally. We define the region between the dorsal (upper third) and ventral (lower third) domains as the ‘equatorial’ domain of the DV (dorsal-ventral) axis, where *Tbx2* shows strong expression (Fig. 1A,B, yellow box), with a rounded termination of its ventral border of expression. In contrast, *Tbx3* displays a more dorsally confined expression pattern, with a distinct spot of enrichment in the equatorial domain that is most prominent between E3 and E4 and largely reduced by E8 (Fig. 1A, middle panel, arrowhead). Notably, both *Tbx2* and *Tbx3* also exhibit a nasal–temporal (NT) component to their expression: *Tbx2* shows a rounded termination at both nasal and temporal borders, whereas *Tbx3* has a straight border that runs diagonally from the nasal and temporal sides towards the enriched spot in the equatorial domain. *Tbx5* is the most dorsally restricted of the three genes, exhibiting a sharp boundary with the equatorial domain (Fig. 1B, yellow box) that remains clearly defined through E8 (Fig. 1A, lower panel). These patterns thus show nested domains of expression along the DV axis, with *Tbx2* furthest ventrally, *Tbx3* enriched in a unique central spot, and *Tbx5* the most dorsally restricted (Fig. 1B).

To assess whether the central spot of *Tbx3* enrichment overlaps with the developing HAA of the chick retina, a multiplexed smFISH approach was used to simultaneously visualize *Tbx3* and *Fgf8* expression. A central spot of high *Fgf8* expression within the equatorial domain serves as a molecular marker of the developing chick HAA , with Fgf8 playing a crucial role in its formation (8, 24). The central spot of *Tbx3* expression was seen to coincide with the spot of high *Fgf8* expression, but not with the broader stripe of Fgf8 expression, demonstrating that *Tbx3* is selectively enriched in the prospective HAA (Fig. 1C).

### Tbx2, Tbx3, and Tbx5 cross-regulate during chick retinal development

To begin to probe the function(s) of the Tbx genes, we performed OE experiments by *in ovo* electroporation of plasmids encoding the Tbx genes. They were delivered at the beginning of neurogenesis, at E2 (Hamburger-Hamilton (HH) stages 9 – 12). Expression was driven by the broadly active chicken beta-actin (CAG) promoter. OE was visualized by use of a ZsGreen reporter gene which was expressed in a polycistronic fashion with the Tbx genes using an IRES sequence. To maintain expression following multiple cell divisions, these plasmids included the sequences necessary for piggyBac transposition, and a piggyBac plasmid was co-electroporated (25) (Fig. 2A). Gene expression in retinal flatmounts was examined at E5 using smFISH (Fig. 2B). In all *Tbx2, Tbx3*, and *Tbx5* OE conditions, ZsGreen signal co-localized with a strong Tbx probe signal for the respective Tbx gene (Fig. 2C-E), confirming successful delivery and expression.

**Figure 2:**
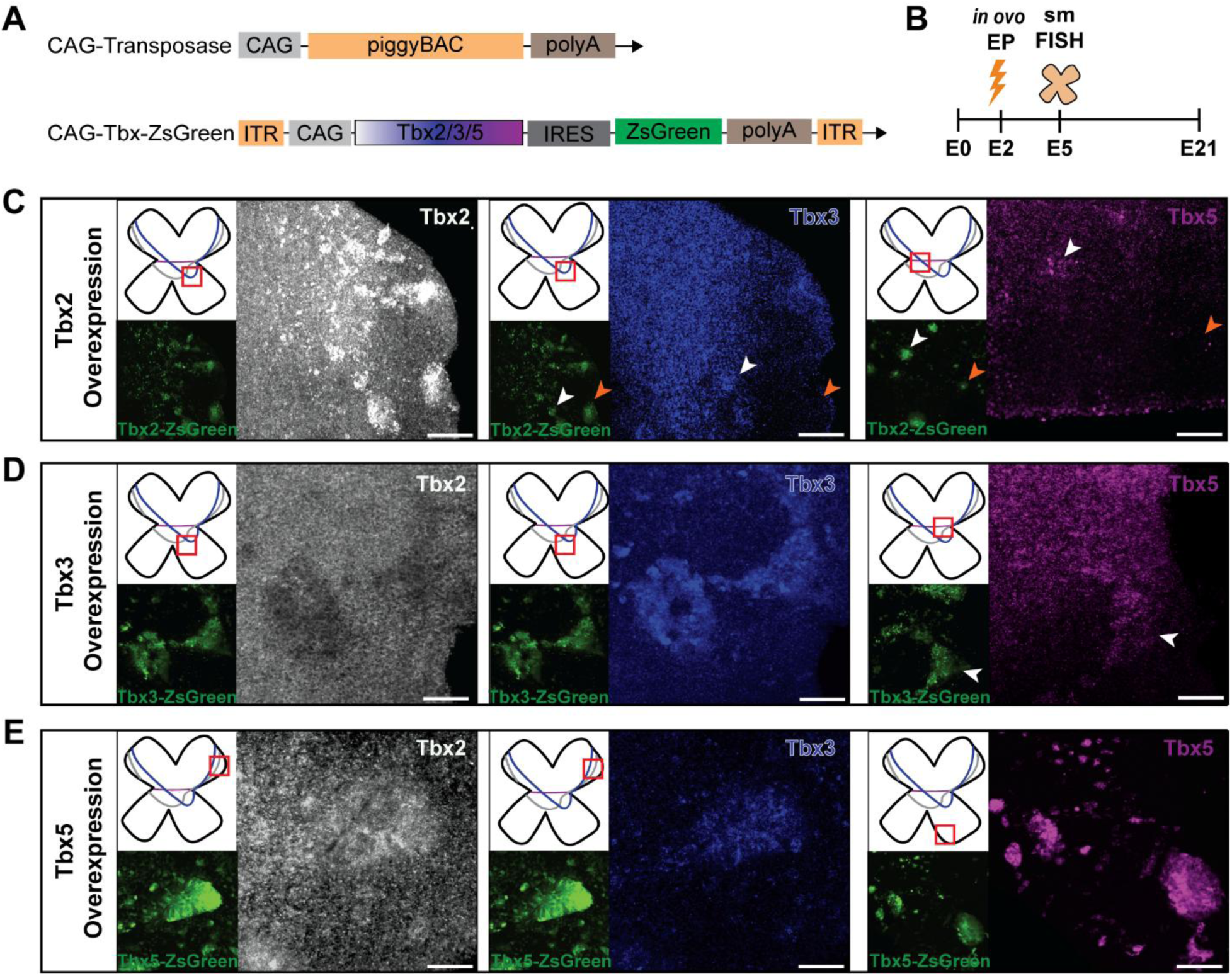
Cross-regulation of Tbx genes in the developing chick retina. **(A)** Diagram of OE plasmids. PiggyBac transposase was used for integration of the Tbx-ZsGreen plasmids. The genes were expressed under the control of the CAG promoter. ZsGreen was used as a fluorescent marker of the electroporated plasmids **(B)** Experimental timeline of the electroporation experiments. **(C-E)** Representative smFISH images of retinal flatmounts electroporated at E2 with Tbx2 **(C)**, Tbx3 **(D)**, and Tbx5 **(E)** OE plasmids and harvested at E5. ZsGreen marks the area of electroporation. Probes for the Tbx genes are indicated in text within each panel. The red square in each panel marks the retinal position at which the representative image was taken. White arrowheads indicate sites of Tbx upregulation within, or adjacent to, the endogenous Tbx expression domain, whereas orange arrowheads mark regions outside the domain where no effect on Tbx expression was observed. Scale bar, 50 µm.

All retinas electroporated with Tbx OE plasmids showed an unusual arrangement of ZsGreen-labelled cells as they appeared in tight clusters. If mCherry, rather than a Tbx gene, was upstream of IRES-ZsGreen, the electroporated cells appeared fairly uniformly distributed in the electroporated region (Fig. S1A). This observation may reflect the activity of Tbx genes on adhesion, migration, and/or survival as reported previously for Tbx genes in other tissues (26, 27). Nonetheless, these retinas were analyzed for their effects on gene expression, and the results were further interpreted in light of results from Tbx knockdown (KD) experiments, as shown later in this study.

OE of *Tbx2* led to a modest upregulation of *Tbx3* in the equatorial retinal domain near the *Tbx3*- enriched spot (Fig. 2C), but not in other retinal domains (Fig. S1B). *Tbx2* OE also resulted in increased *Tbx5* expression within the endogenous dorsal *Tbx5* expression domain, but not outside of it (Fig. 2C and Fig. S1B and S1C).

OE of *Tbx3* resulted in a marked repression of Tbx2 across the entire endogenous *Tbx2* expression domain (Fig. 2D, Fig. S1D). In contrast, *Tbx5* expression was increased both within its endogenous dorsal domain and, to a lesser extent, in the equatorial retinal domain (Fig. 2D, magenta), but not in other regions (Fig. S1D). These findings suggest that *Tbx3* acts as a negative regulator of *Tbx2* while positively influencing *Tbx5* expression.

Notably, *Tbx5* OE induced both *Tbx2* and *Tbx3* expression (Fig. 2E, Fig. S1E). *Tbx2* was strongly upregulated within its endogenous dorsal domain and ectopically in the dorsal periphery, but not in the ventral retina, whereas *Tbx3* was robustly induced throughout the retina, extending into both dorsal and ventral regions. These findings suggest that *Tbx5* acts as a positive regulator of *Tbx2* and *Tbx3*, particularly within the dorsal retina where they are normally co-expressed.

### Analysis of KD of Tbx2, Tbx3, and Tbx5 using mir30-based shRNAs

To determine the requirement of each Tbx gene for the regulation of other Tbx genes, as well as for the expression of other early, patterned genes, knock down (KD) experiments were conducted. A microRNA30 (mir30)-based small hairpin (shRNA) expression system (28, 29) driven by the CAG promoter (CAG-mir30) was used to achieve KD. For stable genomic integration of the electroporated plasmids, the piggyBac transposon system was again used (Fig. 3A) (25).

**Figure 3:**
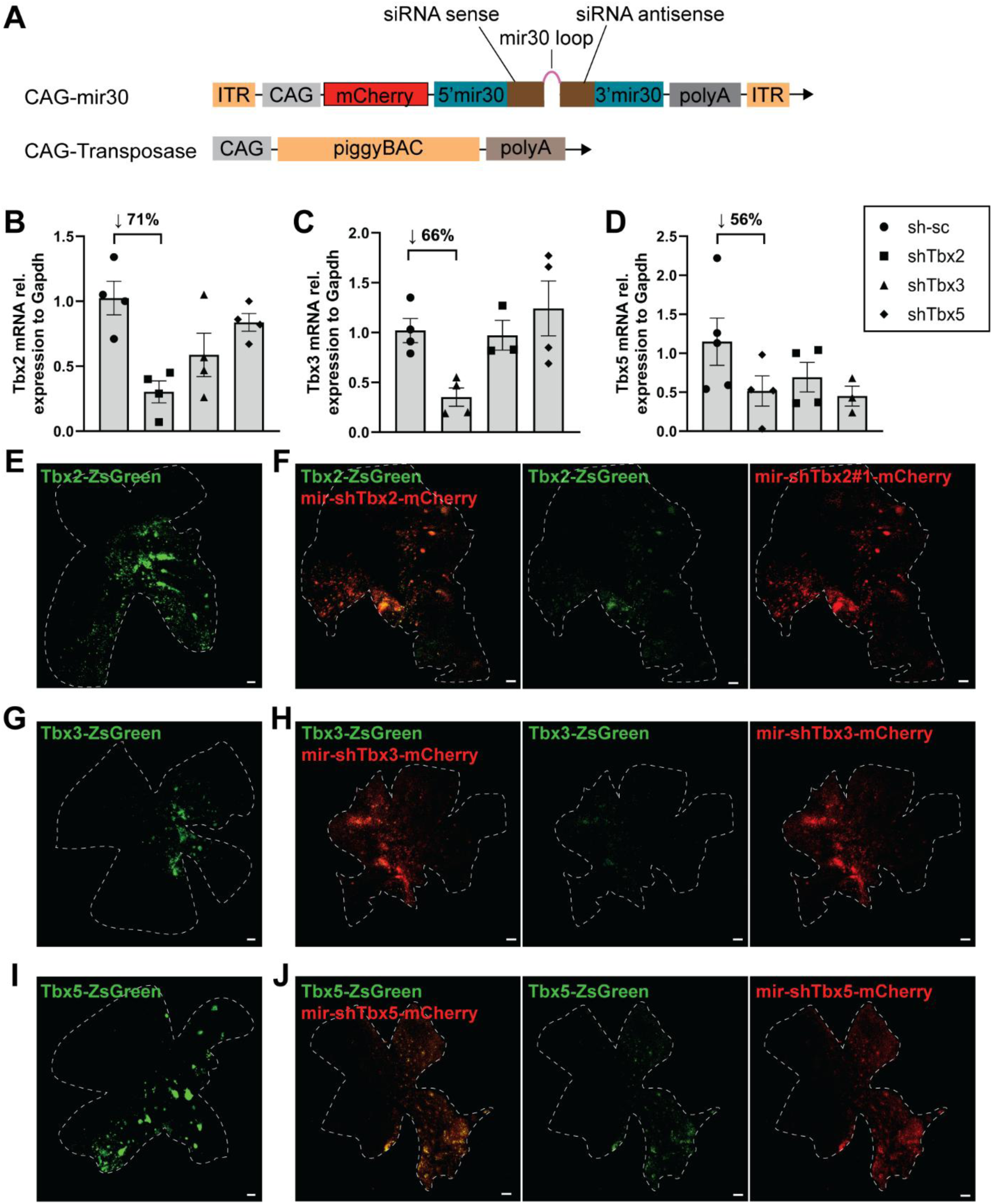
Analysis of efficiency of shRNA-mediated KD of Tbx2, Tbx3, and Tbx5 in vitro and in vivo. **(A)** Diagram of the plasmids for mir30-based RNAi and PiggyBac Transposase. mCherry was used as a marker of the electroporated plasmids **(B-D)** qRT-PCR analysis of KD efficiency for *Tbx2* **(B)**, *Tbx3* **(C)**, and *Tbx5* **(D)** in whole eyecups harvested at E4 after electroporation at E2. Each point represents one biological replicate. Data presented as mean ± SEM. **(E-J)** Representative retinal flatmounts electroporated with a Tbx OE plasmid alone **(E,G,I)** or in combination with the corresponding combination of mir30-shRNA constructs **(F,H,J)**. ZsGreen is expressed from the same transcript as Tbx, serving as an indicator of KD. N = 2 per condition. **(E-J)** Scale bar, 100 µm.

Several KD constructs for each gene were evaluated for efficacy in 293T cells (Fig S2, 25). A non-targeting scrambled shRNA construct expressing mCherry (sh-sc-mCherry) was used as a control for non-specific effects of KD (Fig. S2A-C) (30). KD constructs were also tested for the KD of an unrelated target: OTX2-d2EGFP. The OTX2-d2EGFP signal remained unaffected after co-transfection with any of the Tbx2, Tbx3, or Tbx5 shRNAs (Fig. S2D).

Combinations of shRNAs were used to maximize KD. However, to evaluate specificity, additional shRNA constructs were delivered as single constructs, with a single shRNA used for *Tbx3* and two single shRNAs for *Tbx2* and *Tbx5* (Fig. S3J) These single shRNAs had the same effects on early developmental genes as the combinations of shRNAs (Fig. S3I-M). KD efficiency of the shRNA combinations was quantified by qRT-PCR analysis of whole eyecups collected at E4 after electroporation at E2 (Fig. 3B-D), with efficiencies ranging from 56-71 %. Interestingly, shRNA-mediated KD of *Tbx2* or *Tbx3* also reduced *Tbx5* expression, consistent with our observation that OE of *Tbx2* or *Tbx3* induced *Tbx5* upregulation within its endogenous expression domain (Fig. 2E). In contrast, KD of *Tbx5* did not reduce *Tbx2* and *Tbx3* expression, as might have been predicted from the *Tbx5* OE experiment (Fig. 2E), possibly reflecting supranormal levels of *Tbx5* achieved by OE.

The relatively high variability between biological replicates likely reflects the use of whole eyecups for qRT-PCR analysis. As the size and position of electroporated regions vary considerably among embryos, the proportion of successfully targeted cells differs from sample to sample, thereby reducing the apparent KD efficiency and contributing to inter-sample variability. To address this limitation and further validate KD efficiency in vivo, we co-electroporated the respective Tbx OE constructs either alone or together with the corresponding miR30-shRNA plasmids (Fig. 3E-J). This strategy was chosen because the available smFISH probes for *Tbx2*, *Tbx3*, and *Tbx5* did not reliably detect downregulation of their respective targets, perhaps due to a lack of quantitative read out of the endogenous range of these target RNAs. In addition, antibodies to the Tbx proteins did not reliably work in the tissue. Electroporation of *Tbx2* (Fig. 3E), *Tbx3* (Fig. 3G), or *Tbx5* (Fig. 3I) OE plasmids alone led to strong ZsGreen expression, appearing in bright, patchy domains throughout the retina. In contrast, co-electroporation with the corresponding shRNA constructs resulted in a marked reduction in ZsGreen signal intensity and a more diffuse, less clustered expression pattern across the retina after *Tbx2, Tbx3*, and *Tbx5* KD (Fig. 3F,H,J). Based on both the qRT-PCR and in vivo validation results, it appears that the miR30-shRNA constructs effectively reduced expression of their respective target genes.

### KD of Tbx2/3/5 affects expression of early, patterned genes

Early patterning genes are critical for establishing regional identity within the retina and guiding photoreceptor cell fate (11, 31–34). Additionally, HAA development has been shown to depend on Fgf8 expression and the activity of retinoic acid (RA) degrading enzymes (8). To assess whether *Tbx2/3/5* influence early patterning genes, including those involved in RA metabolism, *in ovo* electroporation was carried out at E2 (HH 8-9) using the mir30-shRNA constructs (Fig. 3). Retinas were collected at E5 and analyzed by smFISH on retinal flatmounts (Fig. 4). Only retinas in which the electroporation marker covered at least 60% of the equatorial and dorsal domains were included in our analysis. To exclude any effects from the electroporation procedure itself and potential off-target effects of the shRNA backbone, all control retinas were electroporated with a mir30-srambled construct (sh-sc-mCherry) at E2 and processed identically. mCherry was used as a fluorescent marker to visualize electroporated cells (Fig. 4).

**Figure 4:**
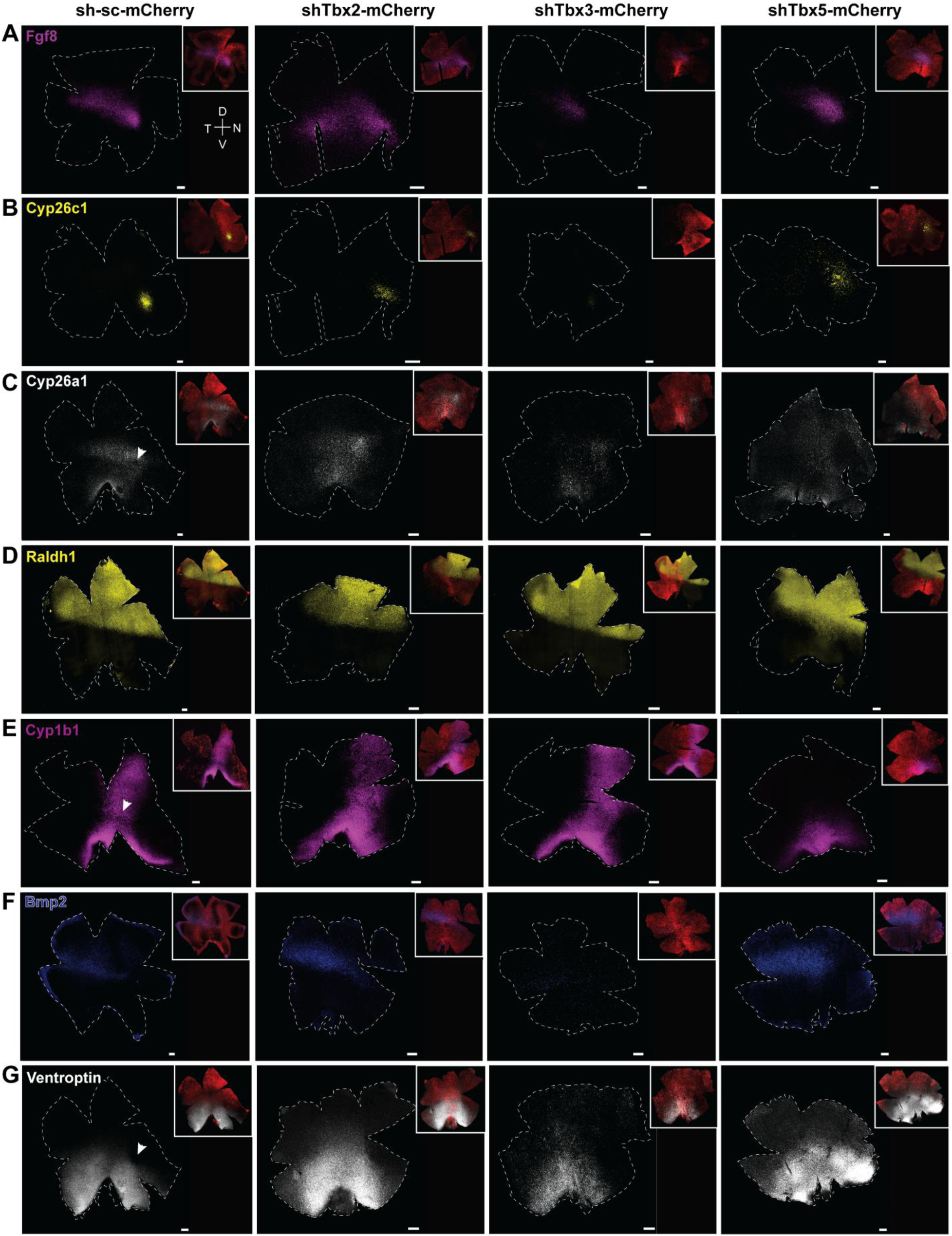
Analysis of effects of KD of Tbx2, Tbx3, and Tbx5 on early retinal genes. **(A-G)** Representative smFISH images of retinal flatmounts electroporated at E2 and harvested at E5. The first column shows control samples electroporated with scrambled shRNA. The electroporated area (mCherry positive) in each representative flatmount is indicated in the upper right corner of each image. The flatmounts were probed for *Fgf8* (Tbx2 KD: N = 3; Tbx3, Tbx5 KD: N = 4) **(A)**, *Cyp26c1* (Tbx2 KD: N = 3; Tbx3, Tbx5 KD: N = 4) **(B)**, *Cyp26a1* (Tbx2, Tbx3, Tbx5 KD: N = 3) **(C)**, *Raldh1* (Tbx2 KD: N = 5; Tbx3 KD: N = 7; Tbx5 KD: N = 3) **(D)**, *Cyp1b1* (Tbx2, Tbx5 KD: N = 4; Tbx3 KD: N = 5) **(E)**, *Bmp2* (Tbx2 KD: N = 4; Tbx3 KD: N = 6; Tbx5 KD: N = 5) **(F)**, and *Ventroptin* (Tbx2, Tbx3 KD: N = 4; Tbx5 KD: N = 3) **(G). (C,E,G)** Arrowheads indicate areas with reduced signal intensity. **(A-G)** Scale bar, 100 µm.

Fgf8 is expressed in the equatorial domain, in a spot and stripe pattern (Fig. 1B, yellow box) (Fig. 4A). As the ventral borders of Tbx expression align with aspects of the Fgf8 pattern, the effects of their KD on *Fgf8* expression were evaluated. KD of *Tbx2, Tbx3,* or *Tbx5* resulted in a complete loss of the defined *Fgf8* pattern, leading to diffuse and reduced *Fgf8* expression (Fig. 4A).

To characterize the influence of Tbx genes on the spatiotemporal expression patterns of RA-degrading enzymes, the expression of *Cyp26c1* (Fig. 4B) and *Cyp26a1* (Fig. 4C) was investigated. Under normal conditions, *Cyp26c1* appears as a distinct, bright spot in the equatorial retinal domain (Fig. 4B), overlapping with the *Fgf8* spot (8). After KD of *Tbx2* with a combination of shRNAs against *Tbx2*, the *Cyp26c1* signal became more dispersed i.e. no longer a definitive spot. An additional single shRNA (i.e. an shRNA not included in the combination of shRNAs used in Fig. 4B) directed against the 3’UTR of *Tbx2* affected *Cyp26c1* (Fig. S3I) and *Fgf8* (Fig. S3J), similarly. KD of *Tbx3* resulted in a smaller *Cyp26c1* domain with reduced signal intensity. Consistent with this, *Tbx3* OE induced ectopic Cyp26c1 expression around the developing HAA (Fig. S3A). In contrast, *Tbx5* KD disrupted the focal expression of *Cyp26c1* in a manner that depended on the size of the electroporated patch. When most of the dorsal retina was electroporated, *Cyp26c1* expression became dispersed and disorganized (Fig. 4B). However, when the electroporated area did not cover the majority of the dorsal retina, *Cyp26c1* expression was largely preserved (N = 4, not shown).

*Cyp26a1* expression also was examined following alterations in Tbx gene expression (Fig. 4C). Under normal conditions, Cyp26a1 is expressed as a broad stripe along the retinal equator, with a distinct but small circular region on the nasal side that does not express Cyp26a1 (8) (Fig. 4C). Additional expression is observed along the optic fissure, with some transcripts present between the fissure and equatorial stripe. Following KD of *Tbx2, Tbx3*, or *Tbx5,* the Cyp26a1-negative spot on the nasal side was no longer visible. Beyond this local change, Tbx KD also broadly altered the overall *Cy26a1* expression pattern. Specifically, *Tbx2* and *Tbx3* KD led to a more diffuse and less defined *Cyp26a1* expression pattern along the equator, while *Tbx5* KD resulted in ectopic expansion of *Cyp26a1* expression into the dorsal domain, extending beyond its normal boundary and appearing more diffuse. Similarly, a single shRNA targeting the *Tbx3* coding sequence caused diffuse expression of *Cyp26a1* (Fig. S3K), consistent with the phenotype observed using two shRNAs in combination (Fig. 4C).

*Raldh1*, a key enzyme in RA biosynthesis, is restricted to the dorsal retina and exhibits a sharp, well-defined boundary at the equator (Fig. 4D). The KD of *Tbx2* or *Tbx3* had no apparent effect on the *Raldh1* expression pattern; the equatorial boundary remained intact, and expression levels appeared unchanged. In contrast, *Tbx5* KD resulted in a blurred *Raldh1* border, suggesting that the equatorial boundary requires intact *Tbx5* activity. This is consistent with the complete overlap of *Tbx5* and *Raldh1* expression domains (Fig. S3B) and is further supported by the observation that *Tbx5* OE induced ectopic *Raldh1* expression throughout the retina (Fig. S3C).

*Cyp1b1*, another enzyme involved in RA metabolism and known to play a role in ocular fissure closure (35) is normally expressed in a vertical stripe extending from the dorsal to the ventral retina (Fig. 4E). While *Tbx3* KD had no discernible effect on *Cyp1b1*, *Tbx2* KD left the overall expression pattern unchanged but caused a modest reduction of expression in the dorsal retina, where the signal appeared dimmer and more diffuse. Strikingly, *Tbx5* KD led to a near-complete loss of *Cyp1b1* expression dorsal to the retinal equator. A single shRNA targeting *Tbx5* was tested and was found sufficient on its own to reduce dorsal *Cyp1b1* expression (Fig. S3L). This observation is supported by the finding that *Tbx5* OE induced ectopic *Cyp1b1* expression, particularly in the ventral retina (Fig. S3D).

The expression of *Ventroptin,* a gene expressed ventrally, and *Bmp2*, expressed in the equatorial domain along the dorsal border of *Ventroptin,* also were examined. Bmp2 plays a key role in retinal development and in maintaining regional specificity along the D/V axis (36). Its expression is antagonistic to that of Ventroptin (36), and both genes exhibit oblique-gradient expression patterns in the developing retina (Fig. 4F,G). *Ventroptin* shows a characteristic depression on the nasal side – precisely coincident with the Fgf8 spot. In contrast to the sharp border around the Fgf8 spot, the dorsal border of the Ventroptin expression domain is diffuse as it meets the equatorial domain (Fig. 3G, S3F) (Joisher et al, BioRxiv). *Tbx2* KD caused a blurred dorsal boundary and dorsal expansion of the *Ventroptin* expression domain (Fig. 4G). This is consistent with *Tbx2* OE, which reduced *Ventroptin* expression (Fig. S3G). Furthermore, *Bmp2* expression appeared dorsally shifted but its overall expression pattern remained intact after *Tbx2* KD (Fig. 4F). Bmp2 may thus act downstream of *Tbx2*, consistent with previous findings (37) alternatively, the expansion of the Ventroptin expression domain dorsally may have caused a dorsal shift in the Bmp2 expression domain. *Tbx3* KD led to a pronounced reduction in *Bmp2* expression, accompanied by dim *Ventroptin* expression that extended dorsally and showed loss of the characteristic nasal depression. In *Tbx5* KD retinas, *Bmp2* expression expanded into the dorsal retina, and *Ventroptin* expression also extended slightly dorsally, with the nasal depression of *Ventroptin* no longer visible. These observations suggest that *Tbx5* functions as a negative regulator of *Bmp2* expression, which is further supported by *Tbx5* OE leading to a reduction in *Bmp2* expression (Fig. S3H).

### Tbx gene KD affects rod photoreceptor patterning and the RFZ

Previous studies have shown that photoreceptor patterning and cell fate decisions rely on spatially restricted gene expression (8, 11, 18, 38). To assess whether *Tbx2, Tbx3,* or *Tbx5* influence photoreceptor distribution and/or fate specification, chick retinas were electroporated at E2 with Tbx shRNA constructs and harvested at E16, after the onset of photoreceptor differentiation (39). Rods and different types of cones were detected using smFISH (Fig. 5A).

**Figure 5:**
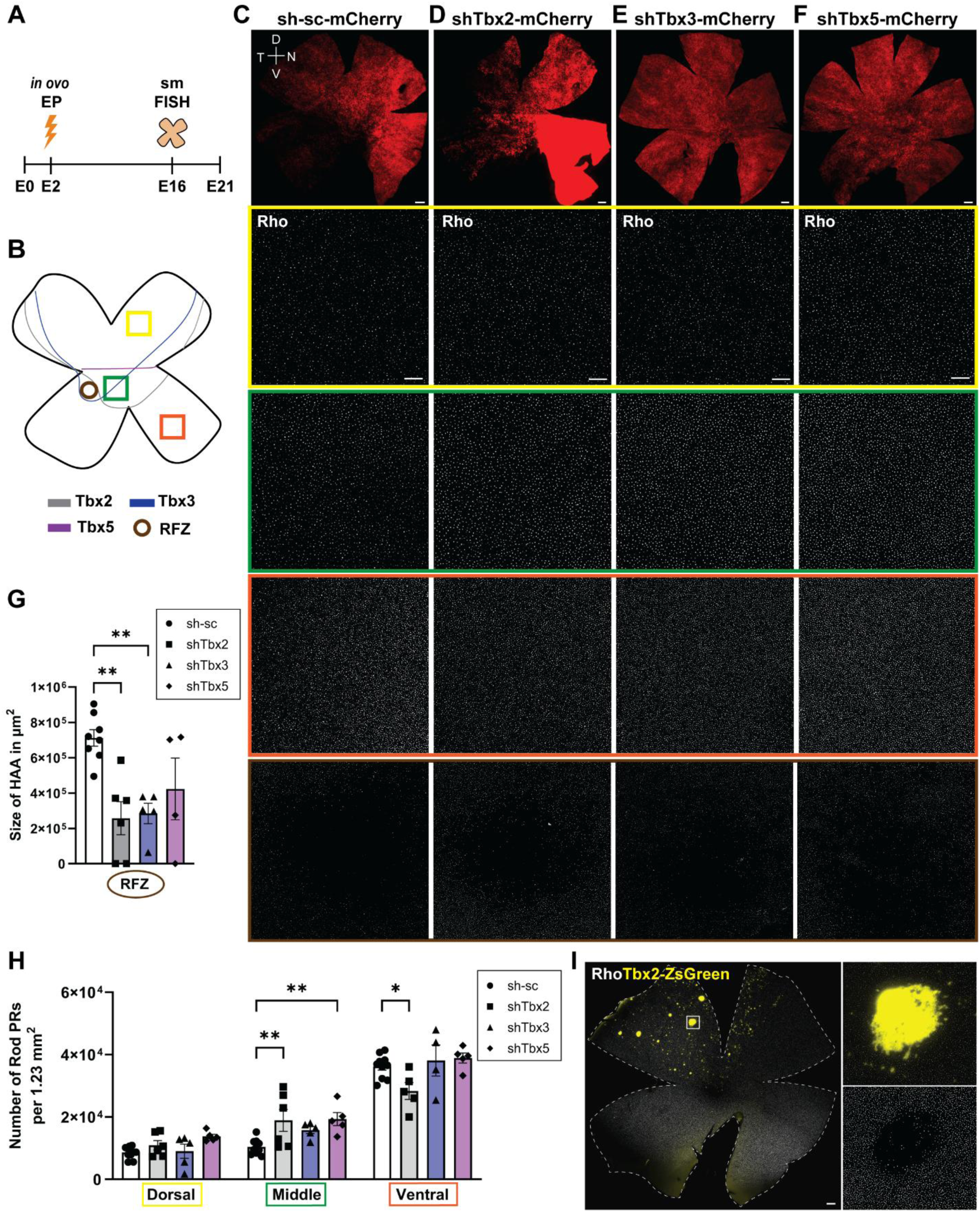
Analysis of rod photoreceptor distribution and RFZ size following Tbx KD. **(A)** Experimental timeline of the electroporation experiments. **(B)** Schematic showing dorsal (highlighted by yellow rectangle), equatorial (green), and ventral (orange) retinal regions analyzed for rod photoreceptor counts. The brown circle marks the RFZ. **(C-F)** Representative smFISH images of E16 retinal flatmounts probed for Rho, following E2 electroporation with sc-shRNA control **(C)**, shTbx2 **(D)**, shTbx3 **(E)**, and shTbx5 **(F)**. Scale bars: mCherry overview images (top row), 500 µm; Rho images, 100 µm. **(G)** Quantification of RFZ size in control and Tbx KD retinas. Control (sh): N = 8; Tbx2, Tbx3 KD: N = 5, Tbx5 KD: N = 4. **(H)** Quantification of rod photoreceptor numbers in dorsal, equatorial and ventral retinal domains. Each point represents one biological replicate. **(G,H)** Data, presented as mean ± SEM, were compared by one-way **(G)** and two-way **(H)** ANOVA. * P ≤ .05, ** P ≤ .01. **(I)** Representative smFISH image of retinal flatmounts electroporated at E2 with a Tbx2 OE plasmid and harvested at E16 and probed for Rho. Scale bar, 500 µm.

Electroporated KD flatmounts were probed for *Rhodopsin* (Rho), a marker of maturing and mature rod photoreceptors (7, 40). Rod numbers were quantified in three retinal regions, allowing a comparison of changes in regional patterning (Fig. 5B). Quantification was restricted to mCherry-positive areas to ensure that only regions with successful plasmid integration were analyzed. Representative images of control flatmounts (scrambled (sc) shRNA; Fig. 5C) and those with KD of *Tbx2* (Fig. 5D), *Tbx3* (Fig. 5E), or *Tbx5* (Fig. 5F) are shown in side-by-side subcolumns to illustrate regional changes in rod numbers. In the dorsal retina, rod numbers were comparable across all conditions, suggesting that KD of these Tbx genes did not impact rod formation dorsally. In contrast, the equatorial retinal domain, located adjacent to the RFZ of the HAA, exhibited a noticeable increase in rod numbers following KD of each of the three Tbx genes. However, this increase reached statistical significance only in *Tbx2*- and *Tbx5*-deficient retinas (Fig. 5H; ANOVA, ** *P* ≤ .01). Consistent with *Tbx2* KD, ectopic OE of *Tbx2* inhibited rod formation (Fig. 5I), reinforcing its inhibitory role in rod photoreceptor development. The ventral retina typically has a higher number of rod photoreceptors than the dorsal region (7). This region remained largely unaffected after *Tbx3* and *Tbx5* KD. KD of *Tbx2* led to a significant reduction in rod numbers in the ventral domain (Fig. 5H; ANOVA, * *P* ≤ .05), while it led to an increase in the equatorial domain. This non-cell-autonomous effect implies that *Tbx2* may influence overall spatial patterning, as well as rod fate specification.

Given the spatially distinct expression patterns of *Tbx2, Tbx3*, and *Tbx5* around the presumptive RFZ and their roles in regulating the expression of patterned genes such as *Fgf8, Cyp26c1,* and *Cyp26a1* (Fig. 4A-C) it was also of interest to analyze the size of the RFZ itself (Fig. 5G). In controls, the RFZ appeared as a well-defined, circular region devoid of rods (Fig. 5C). KD of *Tbx2* and *Tbx3* significantly reduced the size of the RFZ, while its circular shape remained mostly intact (Fig. 5D, E, S4A-C). In contrast, *Tbx5* KD not only reduced the RFZ size but also distorted its shape (Fig. 5F, S4D), making quantification less reliable and resulting in greater variability (Fig. 5G). In some cases, the RFZ was also shifted dorsally (Fig. S4D). Notably, the RFZ was completely absent in two biological replicates after *Tbx2* KD and in one replicate following *Tbx5* KD, further reinforcing the importance of these transcription factors in HAA formation, but also contributing to variability (Fig. 5G). A summary of the diverse effects on RFZ formation after Tbx KD is shown in Fig. S4E.

### Effect of Tbx2, Tbx3, and Tbx5 KD on cone photoreceptors

To further investigate the role of Tbx genes on photoreceptor patterning, cone photoreceptors were quantified following Tbx KD. *Tbx2* KD resulted in a significant reduction of UV cones in the dorsal and ventral retinal domains, as well as in the distorted RFZ, compared to control (sh-sc- mCherry) retinas (Fig. 6A,B). This reduction was statistically significant across all three retinal regions (Fig. 6C; ANOVA, * *P* ≤ .05, *** *P* ≤ .001). In contrast, *Tbx3* and *Tbx5* KD did not noticeably affect UV cone numbers in the dorsal region or the RFZ, but appeared to reduce UV cone numbers in the ventral region, although this decrease did not reach statistical significance (Fig. 6C). The lack of statistical significance and the relatively high variability may in part reflect the fact that the total number of UV cones is approximately six times lower than that of rods, making the data more sensitive to fluctuations between samples. The role of *Tbx2* in UV cone development was further confirmed by *Tbx2* OE experiments, which demonstrated that *Tbx2* OE can induce UV cones (Fig. 6D).

**Figure 6:**
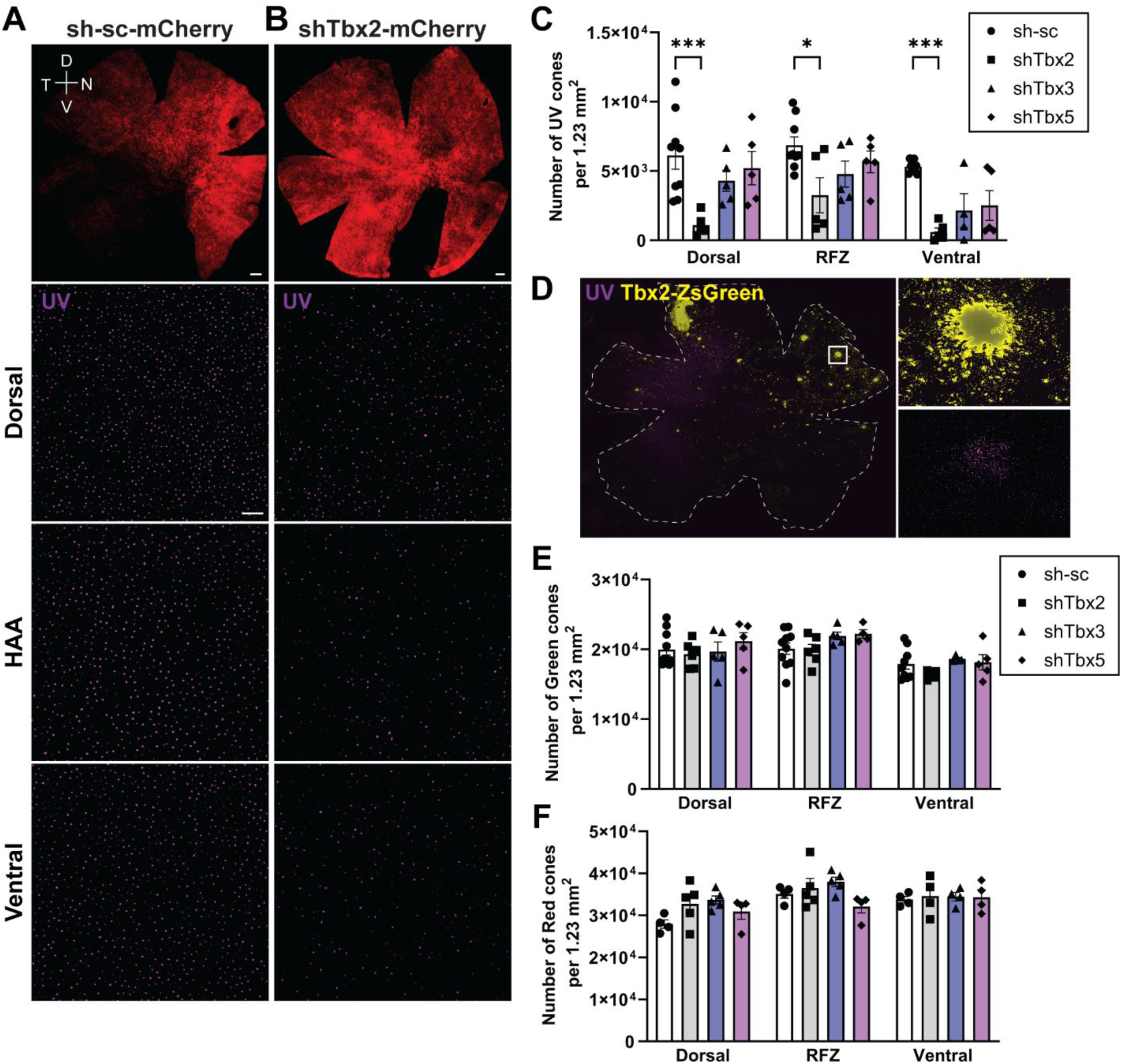
Analysis of Tbx gene alterations on types of cones. **(A,B)** Representative smFISH images of E16 retinal flatmounts probed for UV cones following E2 electroporation with control shRNA-scrambled **(A)** or shTbx2 **(B)**. Scale bars: mCherry overview images (top row), 500 µm; UV cone images, 100 µm. **(C)** Quantification of UV cone numbers in dorsal and ventral domains and in the RFZ in control and Tbx KD retinas. **(D)** Representative smFISH image of a retinal flatmount electroporated at E2 with a Tbx2 OE plasmid, harvested at E16, and probed for UV cones. Scale bar, 500 µm. **(E,F)** Quantification of green **(E)** and red **(F)** cone numbers in dorsal, equatorial, and ventral retinal domains. **(C,E,F)** Each point represents one biological replicate (Fig. S4F). Data, presented as mean ± SEM, were compared by two-way ANOVA. * P ≤ .05, *** P ≤ .001.

The effect of Tbx KD on green and red cones also was examined. In control retinas, green cones were slightly more abundant dorsally than ventrally, and this distribution pattern persisted following *Tbx2*, *Tbx3*, or *Tbx5* KD. Moreover, the total number of green cones in each retinal region remained unchanged compared to controls (Fig. 6E). In control retinas, red cones were more abundant ventrally than dorsally, and this distribution pattern was also not altered by Tbx perturbations (Fig. 6F).

## Discussion

The gain- and loss-of-function experiments reported herein demonstrate that *Tbx2, Tbx3,* and *Tbx5* directly or indirectly regulate each other, while each gene also has its own distinctive effects on the expression patterns of other early genes. They also have some common regulatory roles, as KD of each of them disrupts the early expression patterns of *Fgf8, Cyp26c1,* and *Cyp26a1*. As *Fgf8* and *Cyp26c1* mark the developing HAA, and the RA and Fgf8 pathways play a role in formation of the HAA (8), these results indicate that the Tbx genes also play a role in formation of the HAA. In keeping with this, analysis of a component of the HAA, the RFZ, near the end of development, showed that it was clearly disrupted following KD of each Tbx gene. Rod photoreceptor patterning also was impaired outside of the RFZ in all KDs. *Tbx2* KD additionally led to a smaller number of UV cones. Intriguingly, none of the Tbx KDs affected red and green cones.

### Overlapping expression domains and cross-regulation of Tbx2, 3 and 5

All three Tbx genes are expressed in the dorsal retina, with different ventral borders of expression. OE of *Tbx2* induces *Tbx3* and *Tbx5* within their endogenous dorsal domains, consistent with previous work demonstrating that *Tbx2* is required to maintain the correct spatial expression of *Tbx5* (14). In contrast, *Tbx3* represses *Tbx2*, but induces *Tbx5*, within the normal expression domain of *Tbx5*. This regulatory effect of *Tbx3* on *Tbx2* may indicate a complex balance among Tbx genes, potentially involving shared or partially overlapping functions (41). The upregulation of *Tbx2* and *Tbx3* upon *Tbx5* OE suggests that *Tbx5* serves as central regulator of dorsal retinal identity by coordinating the activation of other T-box members. This creates a mutual activation loop among *Tbx2, Tbx3*, and *Tbx5*, with the notable exception that *Tbx3* represses *Tbx2*, as was reported for cancer cells (42, 43). This inhibitory effect may help refine the expression boundaries within the equatorial region, where *Tbx3* is highly expressed in the presumptive HAA, and *Tbx2* is excluded.

*Tbx2* and *Tbx3* are generally regarded as transcriptional repressors, mediated by their C-terminal repression domains (26, 44). The upregulation of *Tbx5* by OE of either of these genes, only in the normal expression domain of *Tbx5*, suggests that there are other regulators of *Tbx5* that interact with and/or are regulated by *Tbx2* and *Tbx3*. It is also possible that *Tbx2* and *Tbx3* positively regulate *Tbx5* directly as they harbor potential activation domains (13, 42), and they have been shown to positively drive gene expression in a cofactor- and context-dependent manner (37, 42, 45). *Tbx5* is considered to be a transcriptional activator (13, 46), consistent with our findings that it positively regulates *Tbx2* and *Tbx3*.

To examine the individual contributions of these Tbx genes, we designed shRNAs for each gene. Analysis by qRT-PCR revealed that KD of either Tbx2 or Tbx3 led to reduced Tbx5 expression, providing complementary evidence of cross-regulation consistent with the OE experiments. Similar relationships are known from studies of heart development. In the heart, *Tbx2* and *Tbx3* can repress or cooperate with *Tbx5*, in part through interactions with cofactors such as Nkx2.5 and GATA4/6. These transcription factors act at shared or neighboring enhancer elements, and therefore balancing Tbx5-mediated activation and Tbx2/Tbx3-mediated repression to shape the development of the conduction system and chamber myocardium (47–49). In the retina, *Tbx2* and *Tbx3* may similarly stabilize the transcriptional environment required for *Tbx5*, potentially through shared upstream regulatory pathways. The downregulation of *Tbx5* in the *Tbx2* and *Tbx3* shRNA experiments could cause a change in other factors that directly affect *Tbx5* (50, 51). Because of the cross regulatory effects of these Tbx genes, it is possible that any KD effects were due to the loss of more than one family member. This might lead to common effects following KD of each gene, as were seen in some assays, e.g. disruption of the RFZ. However, some aspects of the expression patterns of *Tbx2, Tbx3,* and *Tbx5* did not overlap (Fig. 1A, B), and KD experiments did reveal some specific effects following KD of each Tbx gene, e.g. *Tbx5* KD reduced *Cyp1B1* more strongly than KD of *Tbx2* or *Tbx3*.

### Effects of Tbx OE on electroporated cells

Three days following electroporation of Tbx OE plasmids, the electroporated cells had an unusual appearance. They were arranged in tight clusters, not seen in past experiments or in control electroporated cells in the current studies, e.g. using a plasmid with mCherry upstream of the IRES, rather than a Tbx gene. Tbx OE may be stressful or toxic to retinal progenitor cells. Cells unable to tolerate high levels of these transcription factors may die or exit the cell cycle, leaving behind clusters of “resistant,” ZsGreen-positive cells. Alternatively, the effect may arise from Tbx activities which are not directly toxic. All three Tbx genes are known to influence cell adhesion and migration either directly or indirectly through regulation of genes involved in proliferation and attachment (52–54). For example, Tbx5 OE was shown to prevent proepicardial cell migration (55). In cancer, TGF-β1–induced upregulation of Tbx3 leads to repression of Tbx2, thereby promoting cell migration and invasion (43). Likewise, Tbx3 OE impaired cell growth and migration in skeletal muscle cells (56). These findings collectively suggest that Tbx expression levels must be tightly controlled to maintain normal cellular adhesion, migration, and survival. Despite these caveats, the effects of Tbx OE are likely to be relevant to their gene regulatory roles and photoreceptor patterning, as they complement the KD results with respect to the cross regulation of these Tbx genes.

### KD of Tbx2, Tbx3, and Tbx5 affects expression of early patterned genes

A distinctive pattern in the early chick retina, which we refer to as a "spot and stripe" pattern, was first discovered for Fgf8. In this previous study, we found that Fgf8 is not only patterned in an auspicious manner, but it is required for formation of the HAA, as its KD led to disruption of this area (8). The specific and characteristic Fgf8 expression pattern was disrupted by KD of Tbx2, Tbx3, or Tbx5, indicating that these genes play a role in setting up the HAA. In *Xenopus*, Tbx2 represses Fgf8 signaling by inhibiting Flrt3, a positive regulator of Fgf signaling (57). *Tbx2-* mediated downregulation of Fgf8 could explain the expanded Fgf8 domain we observed following *Tbx2* KD. In contrast, *Tbx5* has been implicated as an indirect regulator of Fgf8 via RA signaling (58). Specifically, in *Xenopus* and mouse cardiopulmonary development, Tbx5 maintains *Raldh2* expression, a RA synthesizing enzyme, thereby antagonizing an Fgf8-Cyp regulatory module to restrict Fgf activity. Consistently, OE of *Tbx5* in the early chick retina showed increased *Raldh1* expression adjacent to the *Fgf8* domain. Although the precise molecular relationship between *Tbx5* and *Raldh1* remains unclear, it has been shown that *Tbx5* expression precedes the onset of *Raldh1* during retinal development (31), further supporting a positive regulatory role. The diffuse *Fgf8* expression after *Tbx5* KD may reflect disrupted RA homeostasis, through disturbed *Raldh1* expression and/or altered expression of RA-degrading enzymes, *Cyp26c1* and *Cyp26a1* Both of these RA-degrading enzymes are likely to shape the *Fgf8* domain (8, 59), and both were suggested to mediate RA degradation within the equatorial mouse retina (60). The observed disruption of the focal *Cyp26c1* expression after *Tbx2* and *Tbx5* KD suggests that *Tbx2* and *Tbx5* are required to establish the spatial boundaries of *Cyp26c1* expression.

High *Tbx3* expression overlaps with the small *Cyp26c1* domain in the developing HAA, whereas *Tbx5* is absent and *Tbx2* is weakly expressed. Our data further suggest that *Tbx3* is required to maintain the size of the *Cyp26c1* domain as well as its expression level, as *Tbx3* KD nearly abolished *Cyp26c1* expression. In addition, *Tbx3* KD resulted in weak and broadly expressed *Cyp26a1*, further pointing to an imbalance in the RA pathway. Interestingly, *Tbx3* KD did not affect *Cyp1b1*, another enzyme that can make RA, whereas *Tbx2* reduced its expression and *Tbx5* abolished it dorsally. Consistently, *Tbx5* OE induced ectopic *Cyp1b1* expression, but only ventrally, suggesting that specific cofactors restrict its transactivation activity dorsally. Together, these findings reveal a previously unrecognized regulatory role of *Tbx2* and *Tbx5* on *Cyp1b1* and further indicate that coordinated regulation of Tbx genes is essential for maintaining the spatiotemporal patterning of RA-degrading enzymes.

We also observed Tbx gene-specific effects on *Bmp2*, a morphogen critical for retinotectal mapping (36). *Bmp2* is expressed from E5 onward in an oblique-gradient pattern complementary to that of *Ventroptin* in the ventral retina (36). *Tbx2* KD did not affect *Bmp2* expression, consistent with prior evidence that *Bmp2* acts upstream of *Tbx2* (36, 37). In contrast, *Tbx3* KD caused strong *Bmp2* downregulation, while *Tbx5* KD led to a dorsal expansion of the *Bmp2* domain. This is consistent with data showing that *Tbx3* can directly regulate *Bmp2* in limb development (61) whereas in heart development, a *Bmp2–Tbx2/3* feed-forward loop operates (62). For *Tbx5*, the data suggest a boundary-setting function: while *Tbx5* expression in the heart is not Bmp2- responsive (63), in the retina *Bmp2* KD reduced *Tbx5* expression (36). Our finding that *Tbx5* KD expanded *Bmp2* expression dorsally, and OE reduced it, supports a boundary-setting function of *Tbx5* for the mid-retinal *Bmp2* domain.

*Ventroptin*, a BMP antagonist, is expressed ventrally and was also affected by Tbx OE and KD. *Tbx2* KD resulted in a dorsal shift of *Bmp2* expression, while *Ventroptin* expanded into the equatorial retina. This is consistent with *Tbx2* acting as a direct *Ventroptin* repressor. This is further supported by our finding that *Tbx2* OE reduced *Ventroptin* expression. This relationship has not previously been reported. *Tbx3* KD did not alter the *Ventroptin* domain size but reduced its intensity and led to reduced *Bmp2* levels. This might be surprising given their antagonistic relationship, perhaps indicating additional levels of regulation in the equatorial domain. Moreover, the loss of the characteristic *Ventroptin* notch marking the *Fgf8* spot following *Tbx3* and *Tbx5* KD correlates with the disrupted *Fgf8* and *Cyp26c1* patterning. Collectively, these findings indicate that *Tbx2, Tbx3*, and *Tbx5* modulate the expression domains of *Bmp2* and *Ventroptin*, potentially allowing aberrant dorsal expansion of Ventroptin - a gene normally confined to the ventral retina - perhaps due to, or thereby causing, disturbed dorsoventral patterning.

Taken together, *Tbx2, Tbx3,* and *Tbx5* KD disrupted domain boundaries separating the equatorial region from dorsal and ventral territories. Interestingly, some of the KD and OE effects occurred in domains where the gene was not expressed, e.g. *Tbx5* OE resulted in ectopic *Raldh1* expression beyond its normal dorsal expression domain and *Tbx2* KD led to reduced ventral rods in a domain larger than the *Tbx2* expression domain. This sort of effect likely reflects aberrant early retinal patterning, which propagate in gene expression changes at a distance.

### Tbx factors regulate photoreceptor cell types and RFZ formation

Perturbations of *Tbx2, Tbx3*, and *Tbx5* resulted in changes in photoreceptor patterning for rods and UV cones, while not affecting red and green cones. KD of *Tbx2* or *Tbx5* resulted in an increased number of rods, but this increase was only significant in the equatorial domain. Neither the dorsal nor the ventral retina showed such an increase, though there was a trend towards an increase in the dorsal region, following *Tbx2* or *Tbx5* KD. In contrast to the increase in rods in the equatorial domain by *Tbx2* KD, the opposite effect was seen in ventral retina, where rods are normally enriched (28). The reduction in rods in the ventral retina occurred in an area outside of the normal domain of *Tbx2* expression. This non-cell-autonomous effect implies that *Tbx2* influences overall spatial patterning, leading to the effect on rods in the ventral retina, and possibly in the equatorial region as well. It may additionally directly affect rod specification in other regions where it is expressed. Supporting this idea, ectopic OE of *Tbx2* inhibited rod formation.

Interestingly, the results of *Tbx2* KD in the ventral retina gave the opposite phenotype to that of Tbx2b and Tbx2a in zebrafish, where the number of rods increased upon loss of Tbx2a or Tbx2b (18), much as we saw in the equatorial domain of the chick retina with *Tbx2* KD. UV cone numbers decreased in the zebrafish Tbx2 mutants, with the numbers of rods and UV cones suggesting a fate switch of UV cones to rods (18, 64). Here we saw a decrease in UV cones in all retinal domains, not just the equatorial domain where rods increased. Even in a control retina, there are almost ten times as many rods as UV cones. Therefore, it would have been difficult to quantify a significant fate switch of UV cones to rods, but this may have occurred in the equatorial domain. The mechanism for the increase in rods in the zebrafish Tbx2 mutants was shown to be due to the lack of repression of Nrl, a gene required for rod formation. The chick has an Nrl homologue, MafA, which may have different regulation from that of zebrafish (65, 66).

As mentioned above, the selective increase in equatorial rods with KD of *Tbx2* and *Tbx5* genes fates may be due to alterations in patterning, through changes in RA signaling. Tbx KD disrupted the balance between RA-synthesizing enzymes (*Raldh1, Cyp1b1*) and RA-degrading enzymes (*Cyp26a1, Cyp26c1*). We previously found that RA injection into chick eyes led to an increase in rods only in the RFZ (8). RA has been shown to promote rod differentiation at the expense of amacrine cells in rat retina (67). In zebrafish, RA exposure shifts fates toward rods and red cones over UV/blue cone types (68). In human retinal organoids, RA is thought to induce formation of green (M) opsin cones and repress formation of red (L) opsin cones (69). Here, we did not detect an effect on the numbers or patterns of red or green cones. M and L cones in chickens likely represent distinct cell types, whereas in humans, M and L cones may represent the same cell type with the only difference being opsin choice, which may be the regulatory step influenced by RA.

In addition to changes in the number of rods in the conditions discussed above, we found that the RFZ was consistently reduced in size and in some cases completely absent following KD of *Tbx2*, *Tbx3*, or *Tbx5*. Formation of all of the features of the HAA – the absence of rods, enrichment of ganglion cells, and a distinctive inner nuclear layer (INL) organization – requires lack of RA and presence of Fgf8 (8). Its formation has not previously been linked to Tbx genes. Although in our experiments we only quantified RFZ size, the consistent reduction (or loss) of the RFZ strongly suggests impaired HAA formation. This suggestion is also supported by the effects of Tbx gene manipulations on the RA and *Fgf8* expression patterns, as RA and Fgf8 perturbations each disrupt all defining HAA features (8).

Taken together, these results demonstrate that *Tbx2, Tbx3*, and *Tbx5* regulate each other, and additionally regulate early retinal genes, including *Fgf8, Cyp26c1, Cyp26a1, Cyp1b1, Raldh1, Bmp2*, and *Ventroptin*. Tbx-dependent positioning of the RA pathway, and potentially of other pathways, are likely at least partially responsible for the downstream effects of *Tbx* KD and OE on photoreceptor fates and the formation of the HAA. These data thus provide a framework to understand not only the molecular mechanisms of HAA development, but more generally how early patterning events influence later cell fate choices.

## Materials and Methods

### Animals

Embryonic retinal tissue was collected from fertilized White Leghorn eggs (Charles River). The eggs were incubated at 38 °C under ∼40 % humidity, and embryos for electroporation were staged according to the criteria of Hamburger and Hamilton (70).

### Tissue preparation

Retinas from the right eye of embryos between E3 - E8 were dissected in 1X PBS without additional treatment, as the retina could be easily separated from the RPE at these stages. For E16 embryos, eyeballs were first cleaned of muscle tissue in 1X PBS and then transferred to a 0.1 mg/mL <u>Dispase</u> I solution (Sigma-Aldrich, D4818; 2 mg dissolved in 20 ml distilled water). The lens and choroid were removed, followed by separation of the RPE. Once detached, the retina was immediately transferred to 1X PBS. Retinas were fixed for 45 min in 4 % paraformaldehyde (v/v) (Thermo Scientific, 28906) at room temperature (RT), then washed 3 × 5 min in 1X PBS prior to HCR RNA-FISH.

### HCR smFISH on retinal flatmounts

HCR smFISH was performed as previously described (24, 71) with minor modifications. Probes were synthesized by Molecular Instruments based on the following sequences: Tbx2 (XM_001235320.7), Tbx3 (NM_001270878.2), Tbx5 (NM_204173.1), Fgf8 (NM_001012767.2), Cyp26c1 (XM_040675448.1), Cyp26a1 (XM_015283751.4), Cyp1b1 (XM_015283751.4), Raldh1 (NM_204577.5), Bmp2 (NM_001398170.1), Ventroptin (NM_204171.2), Opn1sw (NM_205438.1), Opn1msw (NM_205490.1), Opn1lw (NM_205440.2), and Rho (NM_001397497.1). Fixed retinas were washed once in 1X PBS at RT for 5 min, permeabilized in 70 % ethanol/PBS for 2 - 4 h at RT, washed in washing buffer for 5 min at RT, and incubated in hybridization buffer at 37 °C for 30 min. Samples were then hybridized overnight at 37 °C with probes diluted in hybridization buffer (6 µl/ml, 6nM). The next day, retinas were washed twice in prewarmed washing buffer for 30 min each at RT, followed by two washes in 5X SSC (Invitrogen, 15557044) with 0.1 % Tween-20 for 20 min each at RT. Samples were incubated in amplification buffer for 10 min at RT. Meanwhile, matching amplifiers were heated for 90 s at 95 °C, and cooled at RT for 30 min in the dark. Afterwards amplifiers were diluted in amplification buffer (8 µl/ml, 48nM), and applied to retinal samples for overnight incubation at RT in the dark. On the following day, retinas were washed twice in 5× SSCT with 0.1% Tween-20 for 30 min each at RT, then mounted and imaged.

For multiplexing, coverslips were removed in 1X PBS after flatmount imaging. Demounted retinas were incubated with DNase solution (Roche, 04716728001; 30 µl DNase and 150 µl 10X DNase buffer in 1320 µl distilled water) for 1 h at 37 °C. The solution was then replaced with fresh DNase, and samples were incubated overnight at 37 °C. On the following day, retinas were washed twice in wash solution (1 ml 20X SSC and 6.5 ml formamide in 2.5 ml distilled water) for 30 min each at 37 °C, followed by two 30 min washes in 5X SSCT for 30 min each at RT. A second round of smFISH was then performed as described above.

### *In vivo* electroporation

For OE experiments, HH10 - HH11 chick embryos were electroporated in the optic vesicle with CAG-Tbx2-ZsGreen, CAG-Tbx3-ZsGreen or CAG-Tbx5-ZsGreen plasmids (1 µg/ul, final concentration) together with CAG-piggyBacTransposase (1 μg/μl, final concentration). For KD experiments, HH8 – HH10 chick embryos were electroporated with combinations of shRNAs (4 µg/µl each) targeting the respective Tbx gene. A beveled glass capillary needle filled with the plasmid mixture and 0.1 % Fast Green was used to inject the right optic vesicle. To prevent bacterial growth and prevent dehydration, two drops of 1X PBS supplemented with 1 % penicillin-streptomycin (15140148, Thermo Fisher) were added on top of the embryo. Electrodes (Nepagene, CUY610P1.5-1) were positioned extra-embryonically, with the negative electrode placed next to the left optic vesicle and the positive cathode next to the right. The electrodes were gently pressed against the vitelline membrane, and electroporation was performed using a NEPA21 type II electroporater (Nepagene) with five transfer pulses of 15 V, each lasting 50 ms with a 950 ms interpulse interval. Eggs were then sealed with tape and returned to a humidified incubator until the desired developmental stages was reached.

### 293T cell transfection

To identify shRNAs with effective KD efficiency, three candidate shRNAs per gene were tested in HEK293T cells (30). HEK293T cells were cultured at 37 °C in DMEM (11995040, Thermo Fisher) supplemented with 10 % FBS (A4736301,Thermo Fisher), and 1 % penicillin-streptomycin (15140148, Thermo Fisher). Cells at ∼ 70 % confluency in 60-mm dishes were either transfected with a Tbx-overexpressing plasmid with IRES-d2EGFP (unstable GFP) reporter or co-transfected with a CAG-driven mir30-shRNA construct expressing mCherry as an electroporation control. Fluorescent images were acquired 48 h post-transfection using a LEICA DMI3000 B fluorescence microscope.

A combination of shRNAs #1, #2, and #3 for Tbx2 and Tbx5 KD, and shRNAs #1 and #2 for Tbx3, were used for *in ovo* electroporation experiments, with the following sequences: *scrambled (sc)* 5’- CCTAAGGTTAAGTCGCCCTCGTA-3’, *Tbx2#1* 5’-AGCAAATGTCCCATGCCCATA-3’, *Tbx2#2* 5’- TTCTGCAGTTCGTTGAGGGAT-3’, *Tbx2#3* 5’- AGAGCTTTGGGACCAATTCCA-3’; *Tbx2-3’UTR* 5’- ACCGCAATGTTTGGAGTCATT-3’ ;*Tbx3#1* 5’- CCTGGCTAAGCCCATTATGGA-3’; *Tbx3#2* 5’- GACCGCATACCAGAATGATAA-3’; *Tbx3#3* 5’- CGGCCCTAAAGTAGACGAGAA-3’; *Tbx5#1* 5’- GGTCACTGGACTCAATCCAAA-3’; *Tbx5#2* 5’- ATGTCCAGGATGCAGAGTAAA-3’; and *Tbx5#3* 5’-ATTTCTCTGCCCACTTCACCT-3’.

### qRT-PCR

RNA was isolated from retinas with the NucleoSpin® RNA kit (Macherey-Nagel) according to the manufacturer’s protocol. 0.1 – 0.3 µg of total RNA was used for cDNA synthesis using the RevertAid First Strand cDNA Synthesis Kit (Thermo Fisher Scientific) following the manufacturer’s instructions. The obtained cDNA was diluted 1:3 with distilled water and used for quantitative real-time PCR using the PowerUp™ SYBR™ Green Master Mix (Thermo Fisher Scientific) on a CFX96™ Real-Time System (C1000Touch™, Bio-Rad). Exon-spanning primer pairs were designed using the UCSC genome browser. They hybridized exclusively to the desired transcript to avoid detection of genomic contamination. The following primers were used: *Tbx2* 5’- ACCAAATCCGGGAGGAGGAT- 3’ (Forward), 5’- TCGTCGGCCGCAACTATATC-3’ (Reverse); *Tbx3* 5’- TCAAGCTTCCCTACAGCACG -3‘ (Forward), 5’- CAGGGTCAGCTGCTTCCTTT-3‘ (Reverse); *Tbx5* 5’- GACCCTGCTGTCCGCAT-3‘ (Forward), 5’- CGAATCCCCCACTGCGT-3‘ (Reverse); *Gapdh* 5’-GGCCATCAATGATCCCTTCATCG-3’ (Forward), 5’-GGCGTGCCCATTGATCACAA -3’ (Reverse). *Gapdh* was used as the reference gene for data normalization. The “double delta Ct” method (ΔΔCt) was used to calculate the relative expression of the target gene.

### Imaging and quantification

All flatmounts from E16, including those used for quantitative analyses, were imaged with a high-throughput slide scanner (Olympus VS200) using a UPlan X Apo 10x/0.4 air objective lens, and photoreceptor numbers and the RFZ size were quantified in Fiji. For each embryo, cone opsin subtypes were counted within three distinct regions (1.23 mm^2^ each): dorsal, nasal, and the RFZ. Rods were quantified in three regions of the same size, located dorsally, nasally, and temporally adjacent to the RFZ. The size of the RFZ was determined by delineating the border along the transition from the rod-sparse to the rod-dense region, such that portions of the rod-sparse area were included in the measurement. Data were plotted using GraphPad Prism 10.3. Normality was tested with the Shapiro-Wilk test. Values are presented as mean ± standard error of mean (SEM). Photoreceptor counts were analyzed by two-way ANOVA, and RFZ size was compared by one-way ANOVA. Statistical significance was set at P < 0.05 (*P < 0.05; **P < 0.01; ***P < 0.001). N values refer to the number of individual animals per condition, as listed in Fig. S4E.

Control smFISH and Tbx OE retinas were imaged with a Yokogawa CSU-W1 (50 μm pinhole size) spinning disk confocal unit attached to a fully motorized Nikon Ti2 inverted microscope equipped with a Nikon motorized stage and an Andor Zyla 4.2 plus sCMOS monochrome camera. A Plan Apo λ 20x/0.8 DIC I and a Plan Fluor 40x/1.3 Oil DIC H/N2 objective lens were used. Data acquisition was performed with Nikon Elements AR 5.02 acquisition software. Z-stacks were acquired using Piezo Z-device, with the shutter closed during axial movements.

KD retinas were imaged using a single point laser scanning confocal microscope (Leica Inverted DMi8) equipped with a motorized stage (ITK) and five spectral confocal detectors (2 HydS, 2 HyDX, 1 HyDR). All images were acquired using a Plan Apo 20x/0.75 air objective.

## CRediT authorship contribution statement

**Monika Ayten:** Writing – review & editing, Writing – original draft, Visualization, Validation, Project administration, Methodology, Investigation, Formal analysis, Data curation, Conceptualization.

**Heer N.V. Joisher:** Resources, Conceptualization, Methodology, Writing – review. **Constance Cepko:** Writing – review & editing, Supervision, Conceptualization.

## Acknowledgements

The images were acquired using Microscopy Resources on the North Quad (MicRoN) core at Harvard Medical School. We thank Paula Montero Llopis, Praju Vikas Anekal, and Adrienne Wells for their assistance with microscopy and helpful discussions. We thank Ryan N. Delgado and Isabella van der Weide for comments on the manuscript.

