## Supplement for "Tbx genes influence early gene expression and photoreceptor patterning in the chick retina": Ayten et al. Supplement.pdf

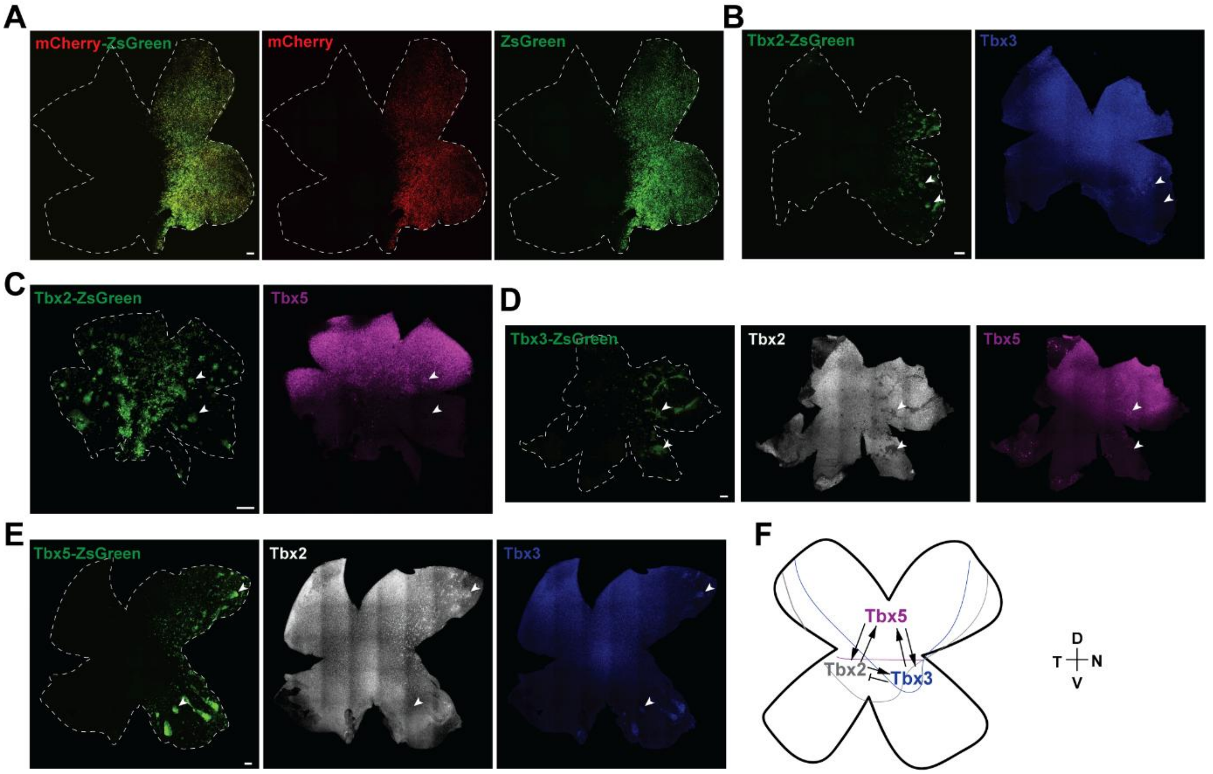


**Figure S1: Mutual regulation of Tbx2, Tbx3, and Tbx5 shapes their expression domains in the embryonic retina (A-D)** Representative smFISH images of electroporated retinas. ZsGreen was used as a fluorescent marker to label electroporated cells. **(A)** Electroporation control. CAG-driven mCherry and ZsGreen. **(B)** Tbx2 overexpression (OE) led to Tbx3 OE in the middle domain of the retina (upper arrowhead), but not in the ventral domain (lower arrowhead). N = 5. **(C)** Tbx2 OE increased Tbx5 expression in the dorsal domain (upper arrowhead) but not outside of it (lower arrowhead). N = 4. **(D)** Tbx3 OE inhibited Tbx2 expression within its endogenous expression domain (arrowheads). N = 4. Tbx3 OE induced Tbx5 expression dorsally (upper arrowhead), to a lesser extent in the middle domain (lower arrowhead), and not at all in the ventral region. N = 3. **(E)** Tbx5 OE induced Tbx2 expression within and outside of its endogenous expression domain in the dorsal region (upper arrowhead), but not in the ventral region (lower arrowhead). N = 2. Conversely, Tbx5 OE induced Tbx3 expression both within and outside its endogenous expression domain, in both the dorsal and ventral regions. N = 4. **(A-E)** Scale bar, 100 µm. **(F)** Schematic representation of the mutual regulation of the T-box transcription factors in the retina.

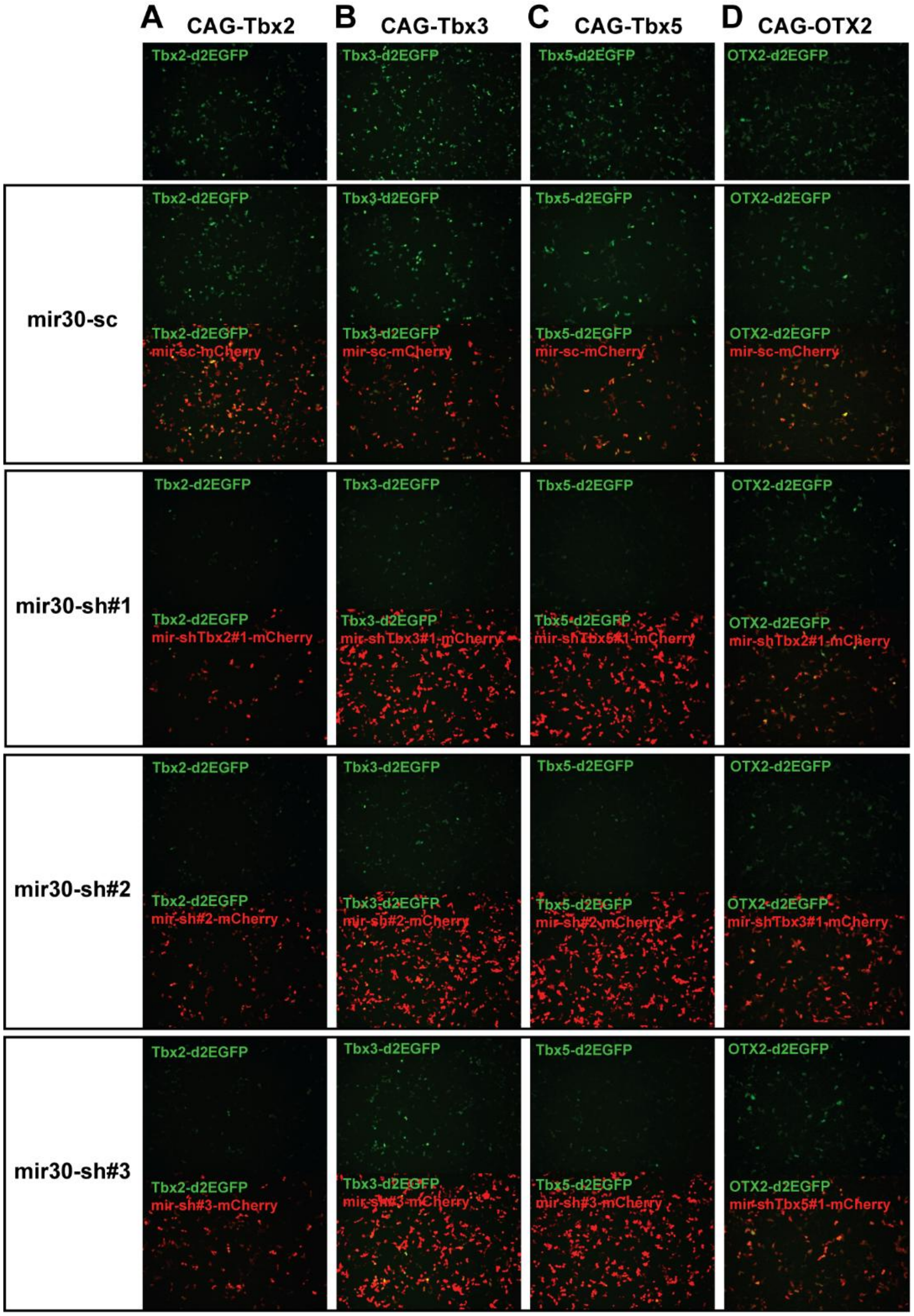


**Figure S2: In vitro characterization of mir30-based shRNA knockdown constructs targeting Tbx2, Tbx3, and Tbx5 in 293T cells. (A-C, top panel)** d2EGFP signal after transfection of Tbx-overexpressing plasmids alone. **(A-C, second panel)** Co-transfection of Tbx-overexpressing plasmids and a scrambled control shRNA (mir30-sc). **(A-C, third to bottom panel)** Co-transfection with Tbx-overexpressing plasmids and gene-specific mir30-based shRNAs targeting Tbx2, Tbx3, or Tbx5 (three shRNAs tested per gene). **(A)** For Tbx2 knockdown: sh#1,#2, and #3 were used in vivo. **(B)** For Tbx3 knockdown: sh#1 and #2 were used in vivo. **(C)** For Tbx5 knockdown: sh#1, #2, and #3 were used in vivo. **(D)** An Otx2 overexpression plasmid was used as an off-target control and transfected either alone, with scrambled shRNA, or with each Tbx-targeting shRNA. None of the shRNAs reduced the Otx2-d2EGFP signal.

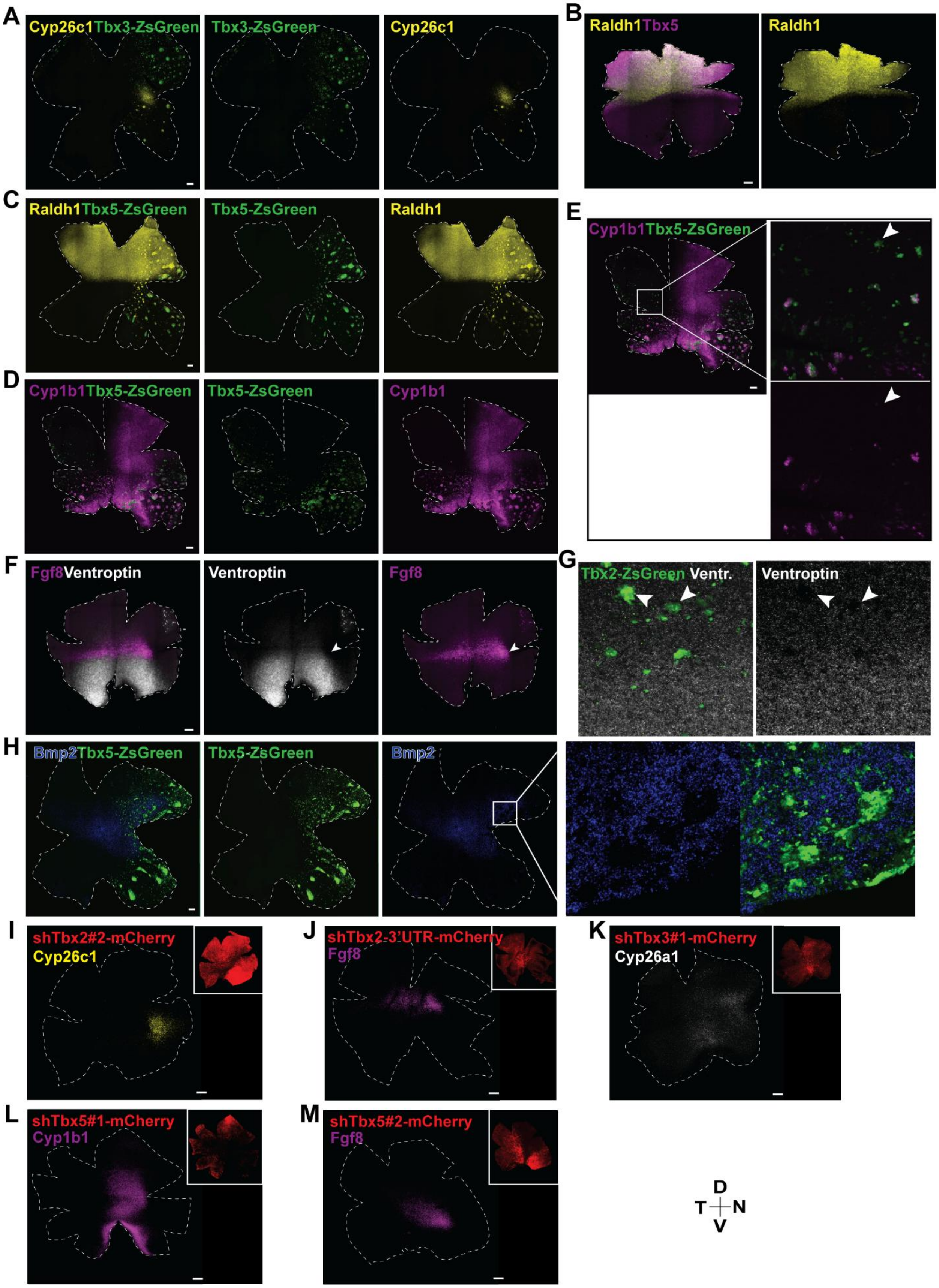

A
Cyp26c1Tbx3-ZsGreen
Tbx3-ZsGreen
Cyp26c1
B
Raldh1Tbx5
Raldh1
C
Raldh1Tbx5-ZsGreen
Tbx5-ZsGreen
Raldh1
D
Cyp1b1Tbx5-ZsGreen
Tbx5-ZsGreen
Cyp1b1
E
Cyp1b1Tbx5-ZsGreen
F
Fgf8Ventroptin
Ventroptin
Fgf8
G
Tbx2-ZsGreen Ventr.
Ventroptin
H
Bmp2Tbx5-ZsGreen
Tbx5-ZsGreen
Bmp2
I
shTbx2#2-mCherry
Cyp26c1
J
shTbx2-3'UTR-mCherry
Fgf8
K
shTbx3#1-mCherry
Cyp26a1
L
shTbx5#1-mCherry
Cyp1b1
M
shTbx5#2-mCherry
Fgf8
D
T
N
V

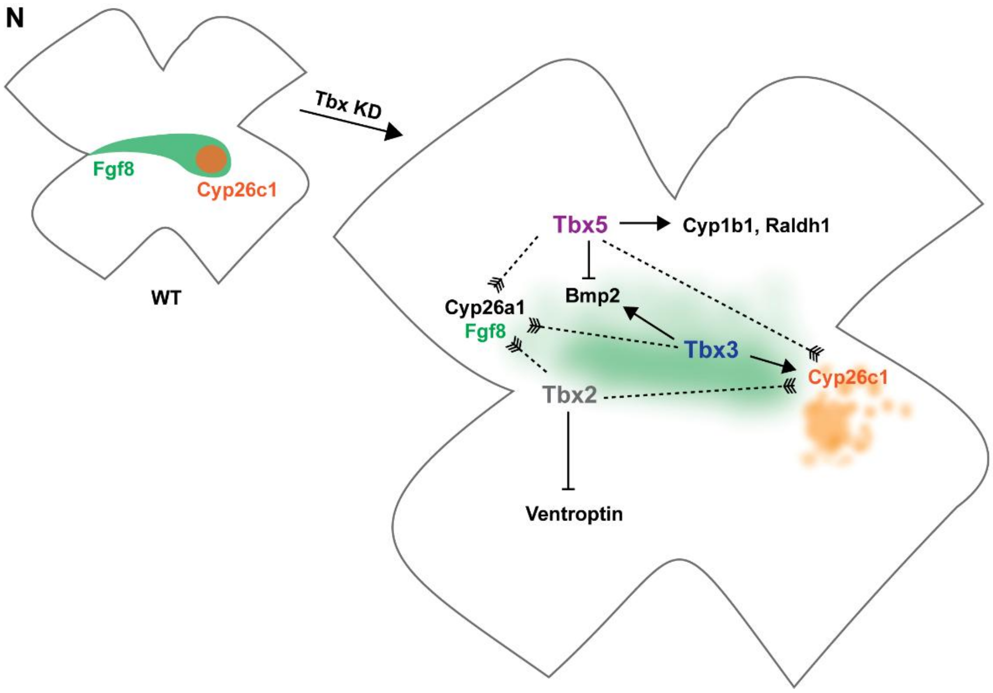


**Figure S3: Complementary electroporation experiments confirm knockdown results. (A,C-E,G)** Representative smFISH images of electroporated retinas. ZsGreen was used as a fluorescent marker to label electroporated cells. **(A)** Tbx3 OE induced ectopic *Cyp26c1* expression adjacent to its endogenous domain in the ventral retina, but not dorsally. N = 4. **(B)** Representative smFISH image of an E5 retinal flamout probed for *Raldh1* and *Tbx5*. **(C)** Tbx5 OE induced ectopic *Raldh1* expression throughout the retina. N = 3. **(D,E)** Tbx5 OE induced *Cyp1b1* exclusively in the ventral retina. N = 3. **(F)** Representative smFISH image of an E5 retinal flatmout probed for *Fgf8* and *Ventroptin*. **(G)** Tbx2 OE repressed *Ventroptin* expression ventrally (arrowheads). N = 3. **(H)** Tbx5 OE repressed *Bmp2* expression and altered its overall distribution, shifting the broader expression domain from the temporal to nasal retina. N = 2. **(I-M)** smFISH images of electroporated retinas with single shRNAs to knock down Tbx2 **(I,J)**, Tbx3 **(K)**, and Tbx5 **(L,M). (A-M)** Scale bar, 100 μm. **(N)** Influence of Tbx2, Tbx3, and Tbx5 on early patterned genes. Arrows indicate genetic relationships i.e. upregulation (arrows), downregulation (flat-headed arrows), and indirect effects on patterning (dashed arrows). Two indirect effects are shown for *Fgf8* (green) and *Cyp26c1* (orange).

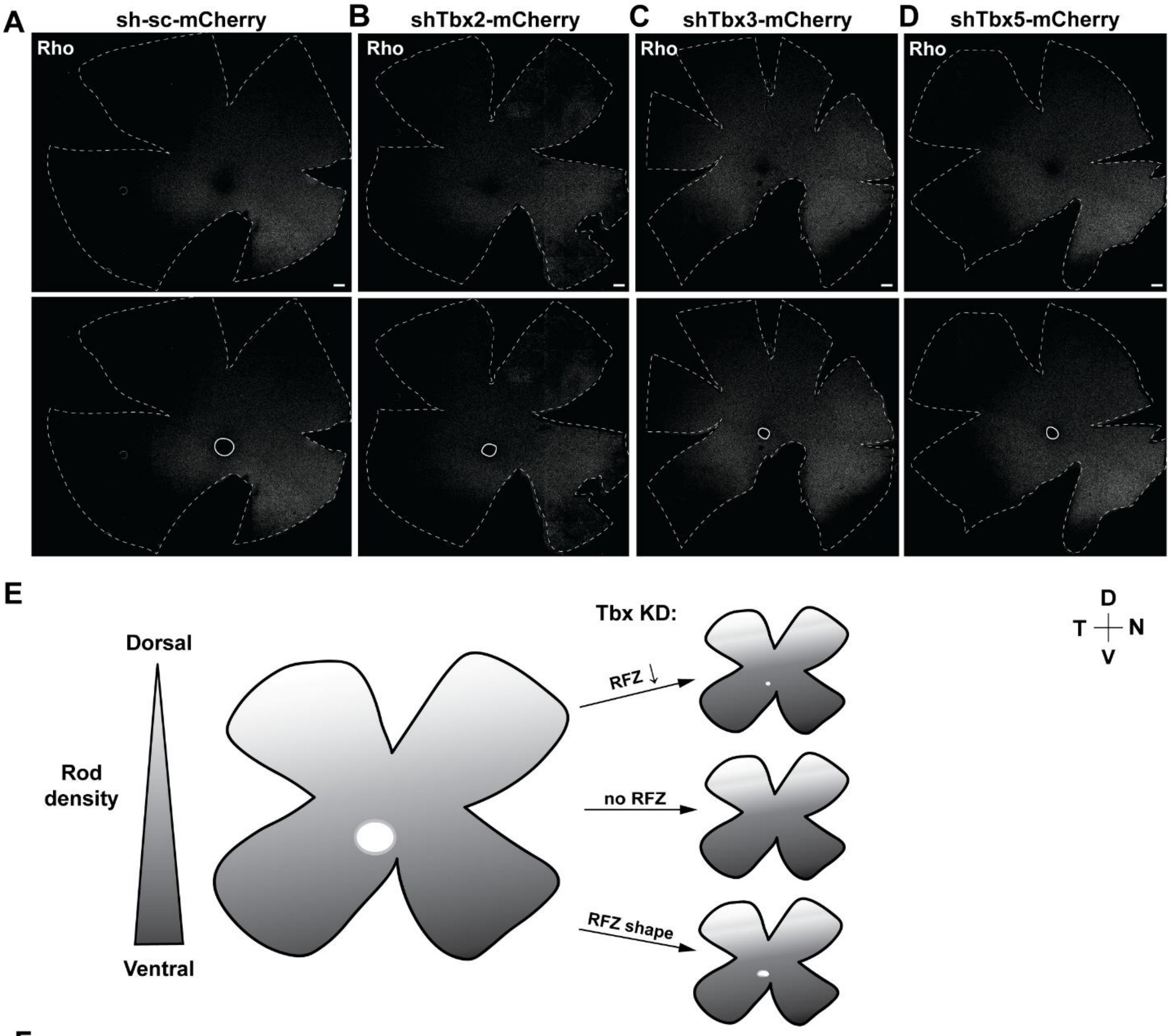


F

| | Rods | | | | UV cones | | | | Green cones | | | | Red cones | | | |
|---|---|---|---|---|---|---|---|---|---|---|---|---|---|---|---|---|
| Knockdown | Control | Tbx2 | Tbx3 | Tbx5 | Control | Tbx2 | Tbx3 | Tbx5 | Control | Tbx2 | Tbx3 | Tbx5 | Control | Tbx2 | Tbx3 | Tbx5 |
| Dorsal | 10 | 6 | 5 | 5 | 9 | 5 | 5 | 5 | 10 | 6 | 5 | 5 | 4 | 5 | 5 | 4 |
| Middle | 10 | 6 | 5 | 5 | 9 | 5 | 5 | 5 | 10 | 6 | 5 | 5 | 4 | 5 | 5 | 4 |
| Ventral | 10 | 5 | 4 | 5 | 9 | 5 | 4 | 5 | 9 | 6 | 4 | 5 | 4 | 4 | 4 | 4 |
| RFZ size | 8 | 6 | 5 | 4 | | | | | | | | | | | | |

**Figure S4: Tbx2, Tbx3, and Tbx5 KD alters RFZ size and positioning. (A–D)** Representative smFISH images of retinas electroporated at E2, collected at E16, and probed for the rod marker *rhodopsin* (Rho). Conditions: scrambled shRNA (**A**), Tbx2 KD **(B)**, Tbx3 KD **(C)**, Tbx5 KD **(D).** Lower panel highlights the RFZ, with Tbx5 KD causing a dorsal shift **(D).** Scale bar, 500 µm. **(E)** Schematic summary of the effects of Tbx KD on RFZ formation, including increased rod density in the equatorial domain. **(F)** Number of biological replicates used for rod, cone, and RFZ size quantifications. Only samples with electroporated coverage of the respective region, strong HCR signal, and intact tissue were considered.
